## Supplemental Figures for "Aging and sperm signals alter DNA break formation and repair in the *C. elegans* germline"

#### Supplemental Figure 1

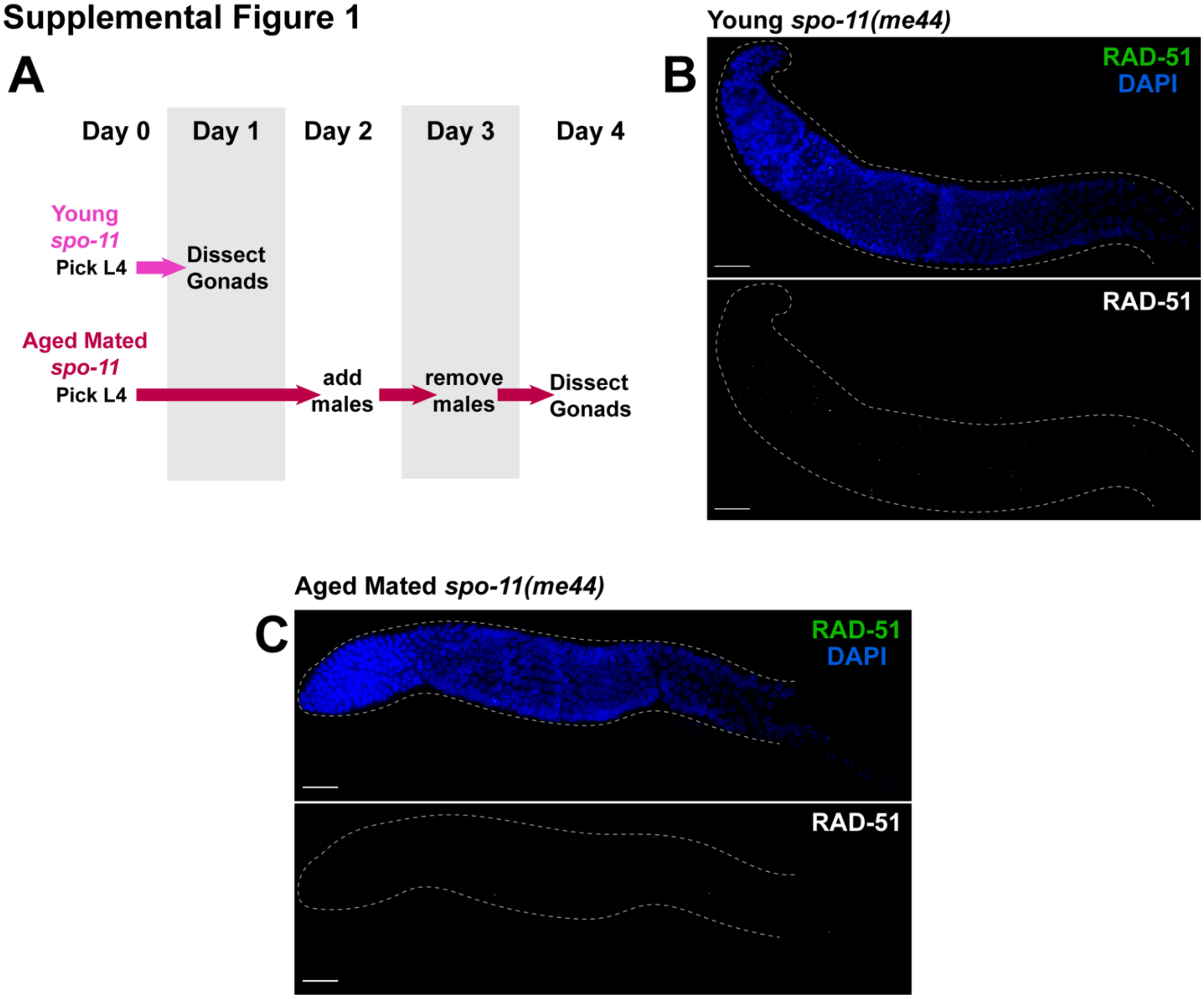


**Supplemental Figure 1. DSBs in aged germlines are SPO-11 dependent.** A) Schemes used to isolate young (1 day post-L4) and aged (4 days post L4) worms for experiments. B-D) Representative whole gonad images of RAD-51 stained germlines from young and aged mated *spo-11(me44)* mutants. Top panels show merged images of both RAD-51 and DAPI, while lower panels show only RAD-51 staining in greyscale. Gonads are oriented with the distal mitotic tip on the left and the end of pachytene on the right. Gonads are outlined with grey dashed lines and scale bars represent 20μm.

### Supplemental Figure 2

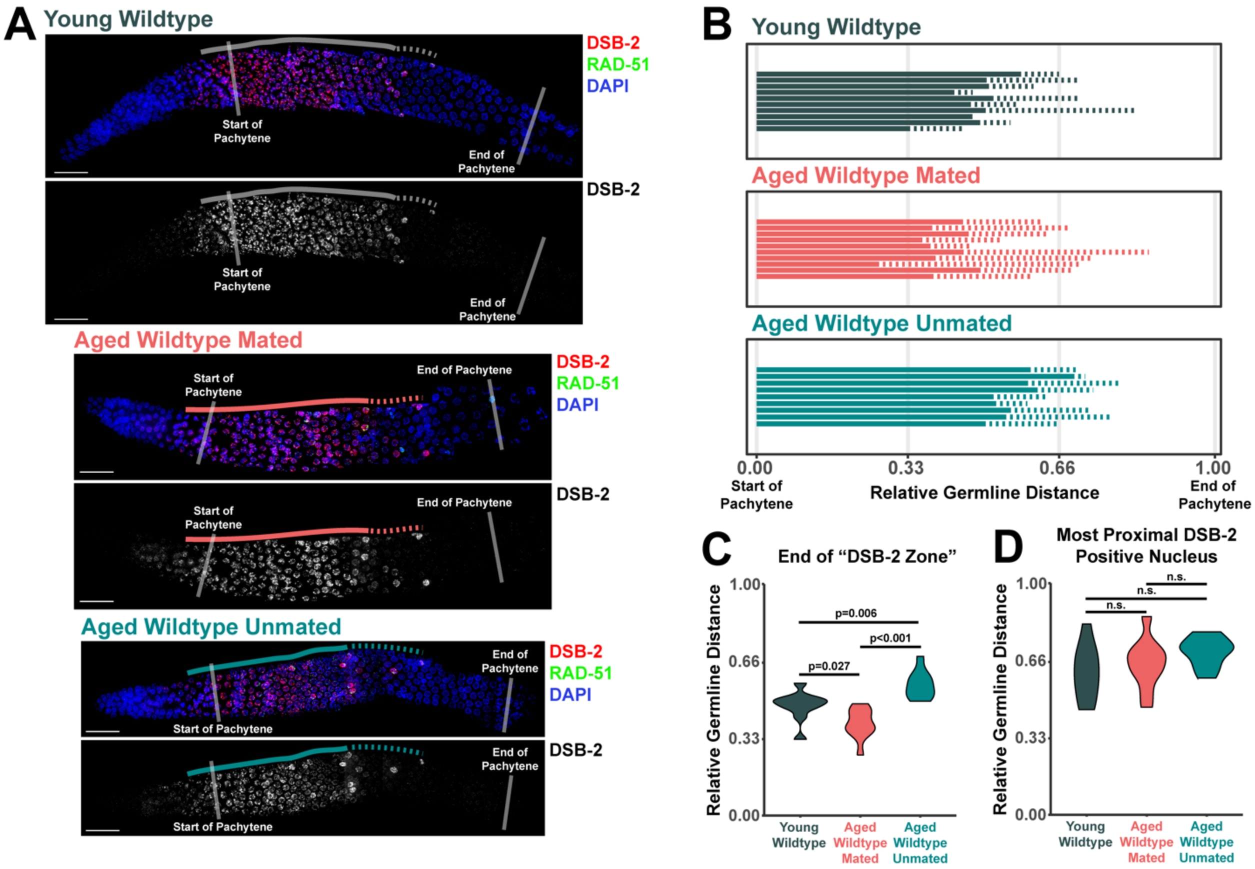


**Supplemental Figure 2. Aged mated and unmated germlines maintain DSB-2 localization in early pachytene.** A) Representative images of germlines stained with DSB-2. Solid lines indicate the "DSB-2 zone", defined as the region of the germline in which >50% of nuclei are stained with DSB-2. Dashed lines extend from the end of the DSB-2 zone to the most proximal nucleus which has DSB-2 staining. Scale bars represent 20µm. B) Line plot representing the quantification of DSB-2 staining in young, aged mated, and aged unmated N2 hermaphrodite germlines. For specific maintenance schemes of these groups, see Figure 1A and Methods. Each horizontal line represents the portion of a single germline which contains DSB-2 positive nuclei. Solid lines represent the "DSB-2 zone", while dashed lines extend to the most proximal germline position at which 1 or more nuclei is marked with DSB-2. C-D) Violin plots comparing the end of the DSB-2 zone and the final position of DSB-2 positive nuclei in young, aged mated, and aged unmated germlines. P values were calculated by Mann-Whitney U test with Bonferroni correction for multiple comparisons.

### Supplemental Figure 3

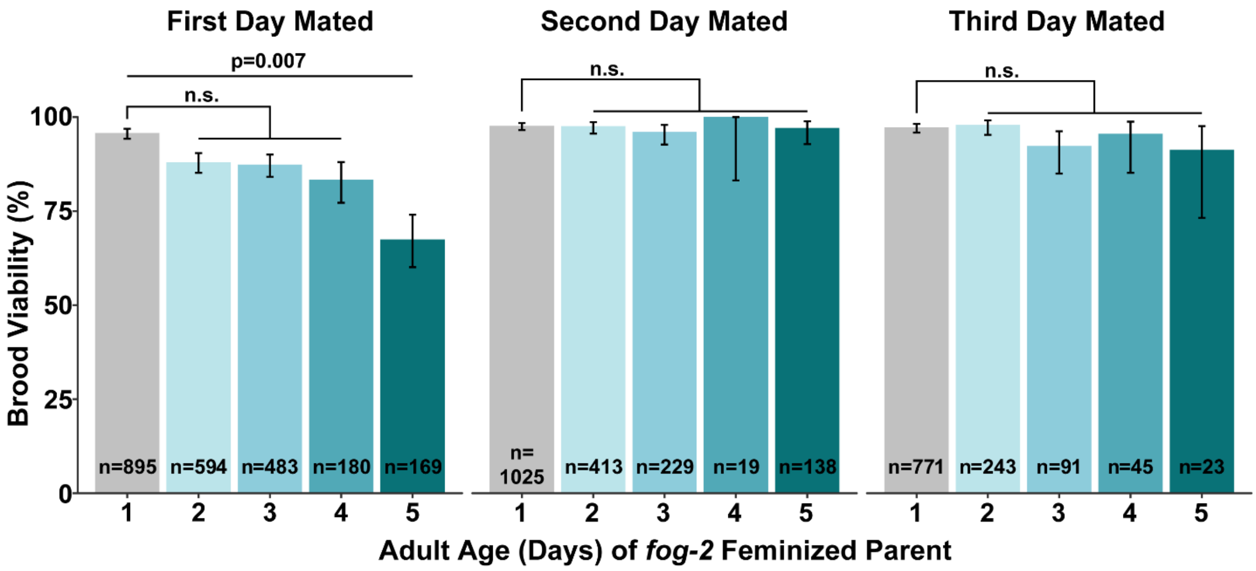


**Supplemental Figure 3. A population of *fog-2* mutant oocytes exhibit reduced viability with parental age.** Bar plots representing the population brood viability of mated *fog-2* mutant females. Error bars represent 95% Binomial confidence intervals. P values were calculated by Fisher's Exact Test. N values indicate the total number of live progeny and dead eggs scored. P values >0.05 are indicated as n.s. (not significant).

#### Supplemental Figure 4

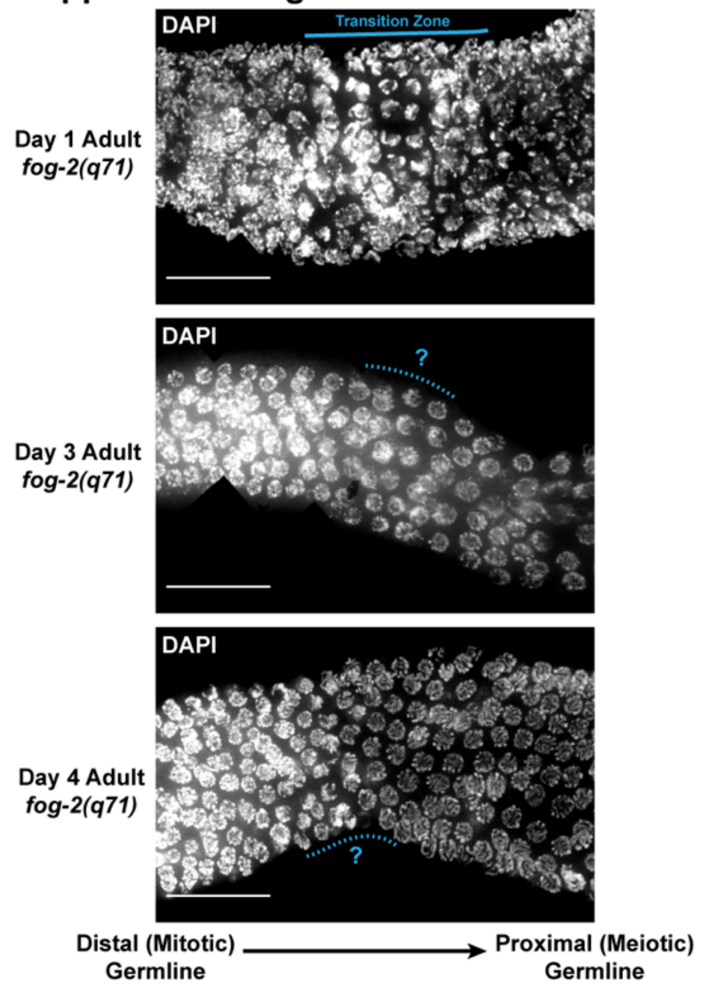


**Supplemental Figure 4. The transition zone is reduced/absent in aged feminized germlines.** Representative images of *fog-2(q71)* feminized mutant germlines from animals 1, 3, or 4 days post-L4. The transition zone is marked with a solid blue line in germlines which have crescent shaped nuclei indicative of meiotic entry. Dashed lines indicate the regions of the germline presumably bridging the mitotic and meiotic germline in which crescent shaped 'transition zone' nuclei are absent. Gonads are oriented with the distal mitotic region on the left and the proximal meiotic region on the right. Scale bars represent 20μm.

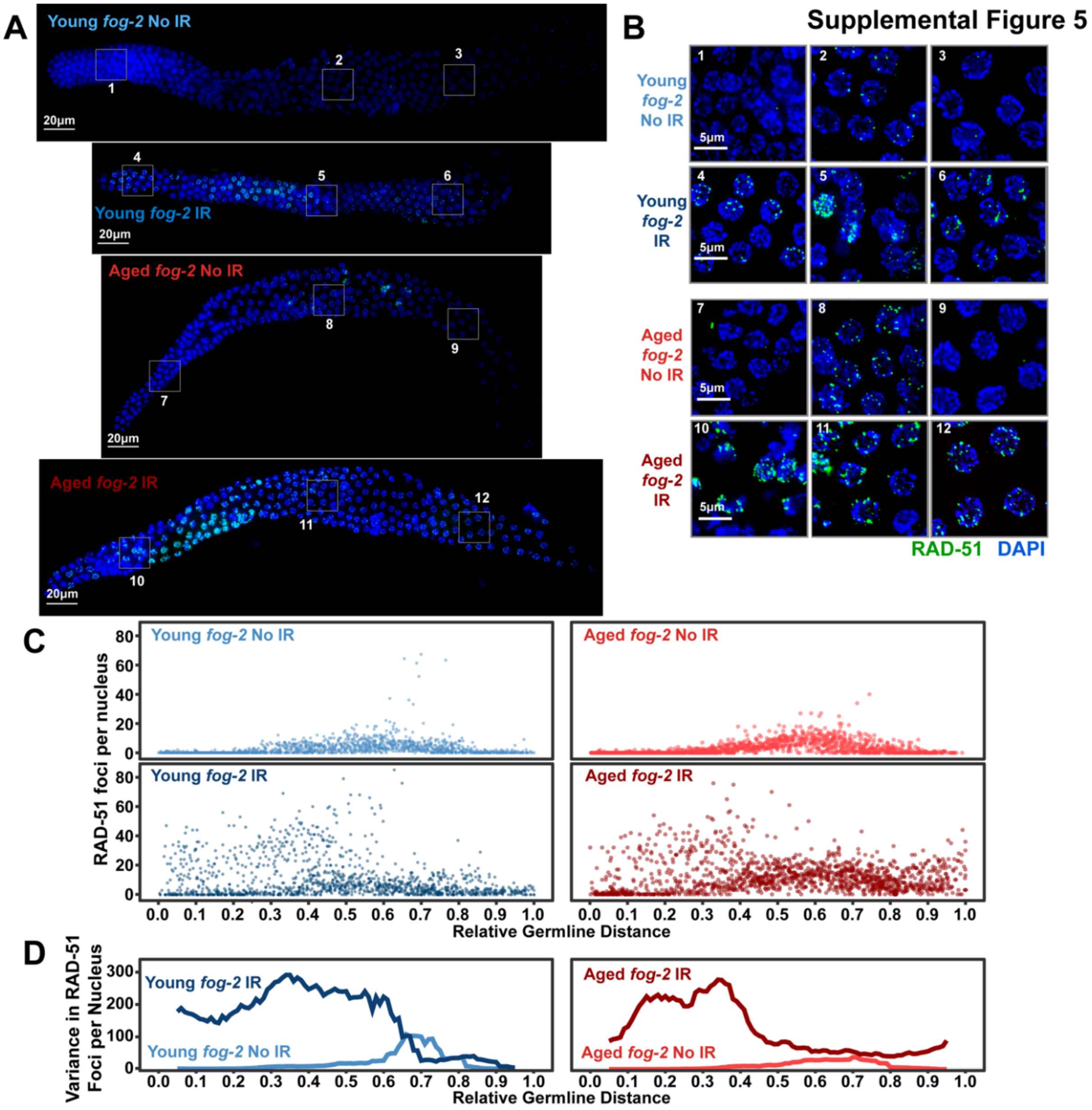


**Supplemental Figure 5. Irradiated *fog-2* germlines exhibit high internuclear variance in DSB repair following irradiation.** A) Representative images of germlines from young and aged irradiated and unirradiated *fog-2(q71)* germlines. For specific maintenance schemes of each group, see Figure 3A. Scale bars represent 20μm. Grey numbered boxes indicate inset panels of nuclei displayed in panel B. B) Represented images of subsets of nuclei from germlines displayed in panel A. Numbers on images correspond to the grey boxes in panel A indicating the portion of the germline each image is derived from. Scale bars represent 5μm. C) Dot plots indicating the RAD-51 foci per nucleus in *fog-2(q71)* IR or No IR young and aged. Each point represents a single nucleus at a given germline position normalized by the premeiotic tip (0) to late pachytene (1) (see Methods). D) Variance in RAD-51 foci per nucleus calculated in a sliding window along the length of the germline where the width of the window is 0.1 germline distance units and the step size is 0.01 germline distance units. Average nuclei quantified in each bin ± standard deviation: Young *fog-2* No IR 141.6±26.6, Young *fog-2* IR 134.4±27.9, Old *fog-2* No IR 148.5±29.6, Old *fog-2* IR 142.4±30.

#### Supplemental Figure 6

***uev-2(gk960600gk429008gk429009)***

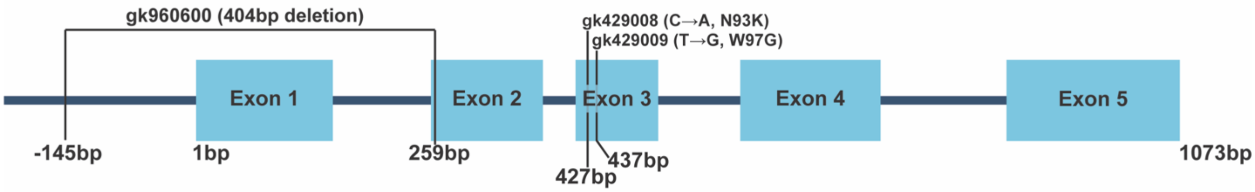


**Supplemental Figure 6. Diagram of *uev-2(gk960600gk429008gk429009)* sequence structure.** Displayed is a scale cartoon of the *uev-2* locus where exons are displayed as boxes and intronic or noncoding upstream sequence is displayed as lines. The gk960600 allele deletes the translation start site and the 5’ intron boundary of exon 2. This lesion generates a frameshift mutation and likely eliminates gene function. Additional point mutations gk429008 and 429009 cause single amino acid substitutions. Base pair distances are indicated relative to the translation start site of Exon 1 of the *uev-2* coding sequence.
